# Designing meta-population genetic management for a small, endangered passerine with fragmented range

**DOI:** 10.64898/2026.02.13.705715

**Authors:** William F. Mitchell, Rebecca Boulton, Rohan H. Clarke, Paul Sunnucks, Alexandra Pavlova

**Affiliations:** La Trobe University, Department of Ecological, Plant, and Animal Sciences, Bundoora, Vic, Australia; The University of Adelaide, School of Biological Sciences, Adelaide, SA, Australia; Monash University, School of Biological Sciences, Clayton, VIC, Australia

**Keywords:** ‘translocation’, ‘conservation’, ‘reintroduction’, ‘Genomics’, ‘genetic rescue’, ‘mallee emu-wren’, ‘Maluridae’

## Abstract

**Context:** Genetic diversity is essential for the persistence and future adaptation of species. However, human-driven habitat fragmentation results in population isolation, often leading to rapid loss of genetic diversity and adaptive capacity. Genetic management of focal taxa may be overlooked in many threatened species conservation programs. The Endangered southeastern Australian mallee emu-wren *Stipiturus mallee* is a species that may benefit from genetic management. Its current range encompasses patchily distributed sub-populations, prone to bottlenecks and genetic drift. Thus, the reintroduction to areas from which the species has been locally extirpated requires careful selection of founders to maximise genetic diversity.

**Aims:** We analyse reduced-representation genomic data from seven sampling areas across the global meta-population to design a translocation strategy that maximises heterozygosity and retention of mallee emu-wren allelic diversity.

**Methods:** We estimated genetic structure, genetic diversity within, and differentiation between subpopulations, thus testing previous inference based on 12 length-variable loci of low population differentiation with 10,840 genome-wide SNP loci. We also estimated effective population sizes to identify populations in need of genetic augmentation, Finally, we used *metapop2* simulations to estimate the relative contributions of each population to global genetic diversity of the species and to estimate the source and number of founders that would maximise heterozygosity and allelic richness in a hypothetical newly established population.

**Key results:** We found weak genetic structure across all sampling areas, supporting previous conclusions that the global mallee emu-wren population should be considered a single genetic unit for management purposes. Low but significant Weir and Cockerham pairwise *F*_ST_ among locations indicated differentiation between sampling areas, suggesting that contemporary gene flow is restricted. Effective population sizes for the two regions supporting the largest numbers of mallee emu-wrens were below the threshold associated with reduced adaptive potential.

**Conclusions:** The genetic health and adaptive potential of sampled mallee emu-wren sub-populations are at risk. **Implications** The global mallee emu-wren meta-population would likely benefit from genetic augmentation, including reciprocal gene flow between extant sub-populations. To maximise genetic diversity in newly established populations, managers should prioritise gene-pool mixing with founders sourced from all sampled areas.

## Introduction

Our planet is in the midst of a human-driven extinction crisis (Ceballos *et al*. 2015). Threatening processes, such as anthropogenic habitat loss and fragmentation, and over-exploitation of natural resources, exacerbated by climate change, have resulted in a hundred-fold increase in global extinction rates when compared with background levels (Ceballos *et al*. 2015; Newbold *et al*. 2015). Small, fragmented populations are susceptible to strong genetic drift and increased likelihood of inbreeding (Courchamp *et al*. 2008; Weeks *et al*. 2016; Frankham *et al*. 2017; Schlaepfer *et al*. 2018). These processes erode genetic diversity (reducing the capacity of a population to adapt to environmental change), reduce fitness (i.e. inbreeding depression), and if left unmanaged, may result in a feedback loop of accelerated population decline, ultimately leading to extinction (Saccheri *et al*. 1998; Keller and Waller 2002).

Despite its significance, maintenance of genetic diversity of focal species has been poorly incorporated into conservation management due to underappreciation of the importance of genetic variation, inertia, and limited knowledge of how to apply evolutionary thinking in conservation (Pierson *et al*. 2016; Cook and Sgro 2017; Laikre *et al*. 2020). In recent years, several researchers have advocated for conservation management that explicitly considers the maintenance of genetic diversity and evolutionary processes at a meta-population scale (Ralls *et al*. 2018; Hoban *et al*. 2021; Pavlova *et al*. 2025; Shaw *et al*. 2025). This objective is often achievable through genetic augmentation, such as the translocation of living organisms between fragmented populations, thus improving genetic health and adaptive potential (Frankham *et al*. 2017; Liddell *et al*. 2021).

Many species form meta-populations with gene flow amongst sub-populations maintained through temporally or spatially dynamic connectivity (Bertassello *et al*. 2021). For example, periods of flooding may temporarily connect otherwise isolated waterbodies while rainfall and fire history drive variable habitat suitability and connectivity for many species in semi-arid and fire-prone terrestrial habitats (Piana *et al*. 2014; Connell *et al*. 2022). Within these dynamic systems, local extinction and re-colonisation events are fundamental processes that drive meta-population dynamics (Hanski 1998). However, anthropogenic impacts (e.g., altered water or fire regimes, or other habitat degradation) can interrupt these processes, leading to isolated populations at greater risk of genetic consequences, and little probability of re-colonisation following local extinctions (Camak *et al*. 2021; Knipler *et al*. 2023). The likelihood of ongoing persistence of such meta-populations may be enhanced through management interventions designed to increase genetic diversity within and across sub-populations (Pavlova *et al*. 2025).

Sub-populations newly re-established by translocation following local extinction are invariably founded from a subset of individuals—and hence a subset of the genetic diversity—from a source population (He *et al*. 2016). This subsampling places an upper limit on the genetic diversity that a population may contain at establishment (Nei *et al*. 1975; Hundertmark and Van Daele 2010; Andersen *et al*. 2014). Newly established populations are also typically small (Langridge *et al*. 2021), rendering them susceptible to the processes described above that may erode genetic diversity and fitness (Andersen *et al*. 2014). For these reasons, translocation managers should seek to optimise genetic diversity through careful selection of founders based on quantitative assessment of genetic richness of potential source populations (Malone *et al*. 2018; Nistelberger *et al*. 2025). Translocations may also be used to augment gene flow for small, isolated populations, providing a source of novel genetic material (Grueber *et al*. 2017). By introducing individuals from larger, more diverse populations, from a greater number of populations (Lutz *et al*. 2021), by selecting founders with high individual heterozygosity (Scott *et al*. 2020; Feuerstein *et al*. 2024), or conducting reciprocal translocations between multiple small populations (Heber *et al*. 2013; Frankham 2015), managers may improve translocation success, alleviate the effects of inbreeding, bolster fitness and improve the adaptive potential of populations.

Genetic data can guide genetic management to improve population fitness and ability to adapt to future environmental changes (Pavlova *et al*. 2025). For example, effective population size, *N*e (the number of individuals reproducing in an ideal population that would result in the same level of genetic drift observed in a real population) could indicate a population’s vulnerability to genetic drift (*N*e < 1000 would flag that a population is at risk of low adaptive potential, Frankham *et al*. 2014). Similarly, increasing differentiation between previously connected populations is a signal of disrupted natural gene flow (Amos *et al*. 2014). In addition, simulation approaches could guide selection of sources for translocation to minimize inbreeding in newly-established populations and maximize retention of useful genetic variation (Rick *et al*. 2023; Pavlova *et al*. 2025).

Here we use genetic analyses to guide genetic management of the global population of the Endangered mallee emu-wren *Stipiturus mallee*. The mallee emu-wren is a small passerine endemic to the semi-arid mallee region of north-western Victoria and eastern South Australia (Higgins *et al*. 2001). The species is now extinct in seven reserves it once occupied, including all South Australian sites. An estimated <7,500 individuals remain, with the majority (∼6,500) distributed patchily in the northern part of their extant range (Verdon *et al*. 2021; Verdon *et al*. 2025). Previous genetic research using twelve length-variable loci (11 microsatellites and one EPIC marker) characterised genetic diversity across mallee emu-wren populations as low to moderate with evidence of genetic bottlenecks in five of six sampling locations (Brown *et al*. 2013). The strongest signal of demographic and genetic impoverishment was observed in the Ngarkat Conservation Park sub-population (Brown *et al*. 2013), which became extinct after 2014 (Mitchell *et al*. 2021). Weak spatial population genetic structure suggested some degree of gene flow among sampling areas, supporting treatment of the whole species as a single genetic unit (Brown *et al*. 2013). Subsequent to that study, an isolated population of mallee emu-wrens was rediscovered in Wyperfeld National Park in 2015, and little is known about the genetic health of this population.

Translocations are identified as a priority conservation action to re-establish mallee emu-wrens in isolated habitat that has recovered following fire, and to provide insurance against loss of extant sub-populations to future wildfire (Indigo *et al*. 2023). Accordingly, the first trial reintroduction of mallee emu-wren occurred in 2018 into Ngarkat Conservation Park in South Australia. That trial did not succeed in reintroducing a population but did improve knowledge of husbandry, translocation protocols and general ecology of the species (Hunt *et al*. 2019; Mitchell *et al*. 2021). A new mallee emu-wren reintroduction is currently under consideration. A more informative understanding of the genetic composition of different potential source populations, allowing managers to maximize offspring fitness in newly translocated populations (by minimizing inbreeding and maximizing retention of unique genetic variation) could support future conservation efforts.

In this study we use a genomic dataset of 10,840 single nucleotide polymorphism (SNP) loci genotyped for 71 individuals, including three from a population in Wyperfeld National Park, to inform genetic management of the mallee emu-wren across its entire distribution. We aim to:

1. Re-examine range-wide genetic structure and genetic differentiation among sampling areas with greater genomic and spatial resolution than a previous study,
2. assess levels of individual heterozygosity and population genetic diversity within sampling areas,
3. estimate effective population size for cohesive genetic populations and identify populations at risk of inbreeding depression that may benefit from gene flow augmentation, and
4. identify optimum sources of founders for future translocation or captive breeding programs that will maximise retention of the species’ genetic diversity.

## Methods

### Study system

Mallee emu-wrens are confined to two contiguous tracts of remnant native vegetation (referred to here as ‘regions’) in south-eastern Australia: Hattah-Kulkyne National Park, Murray Sunset National Park and contiguous reserves (northern mallee region) and Wyperfeld National Park (southern mallee region, Fig. 1, Verdon *et al*. 2021; Verdon *et al*. 2025). The mallee emu-wren prefers dense cover of complex low-lying vegetation, particularly *Triodia scariosa*, and is rarely observed flying (Higgins *et al*. 2001). Population trajectories for this species are dynamic and influenced by prevailing climatic conditions and time since fire, with gene flow more likely during periods of above-average rainfall and within areas of appropriate fire age (Brown *et al*. 2009; Verdon *et al*. 2019; Connell *et al*. 2022; Mitchell *et al*. 2025). Historically, gene flow was likely maintained over long time periods across a large mallee emu-wren meta-population encompassing suitable mallee vegetation south of the Murray River in Victoria and South Australia. Since extensive clearing for agriculture in the early 20^th^ century, mallee emu-wrens have gradually disappeared from those remnant areas that are small, isolated or of marginal quality due to reserve-scale fire, often exacerbated by drought. Natural immigration is not considered feasible across cleared or degraded land, precluding re-colonisation following fire, despite subsequent recovery of suitable habitat. Wildfire poses a great risk in this system. Long-unburnt areas of high-quality habitat have disproportionally high occurrence of mallee emu-wrens (Verdon *et al*. 2019). A reserve-scale wildfire encompassing such an area of core habitat (e.g., Hattah-Kulkyne National Park) could threaten global persistence of the mallee emu-wren (Indigo *et al*. 2023).

**Figure 1.**
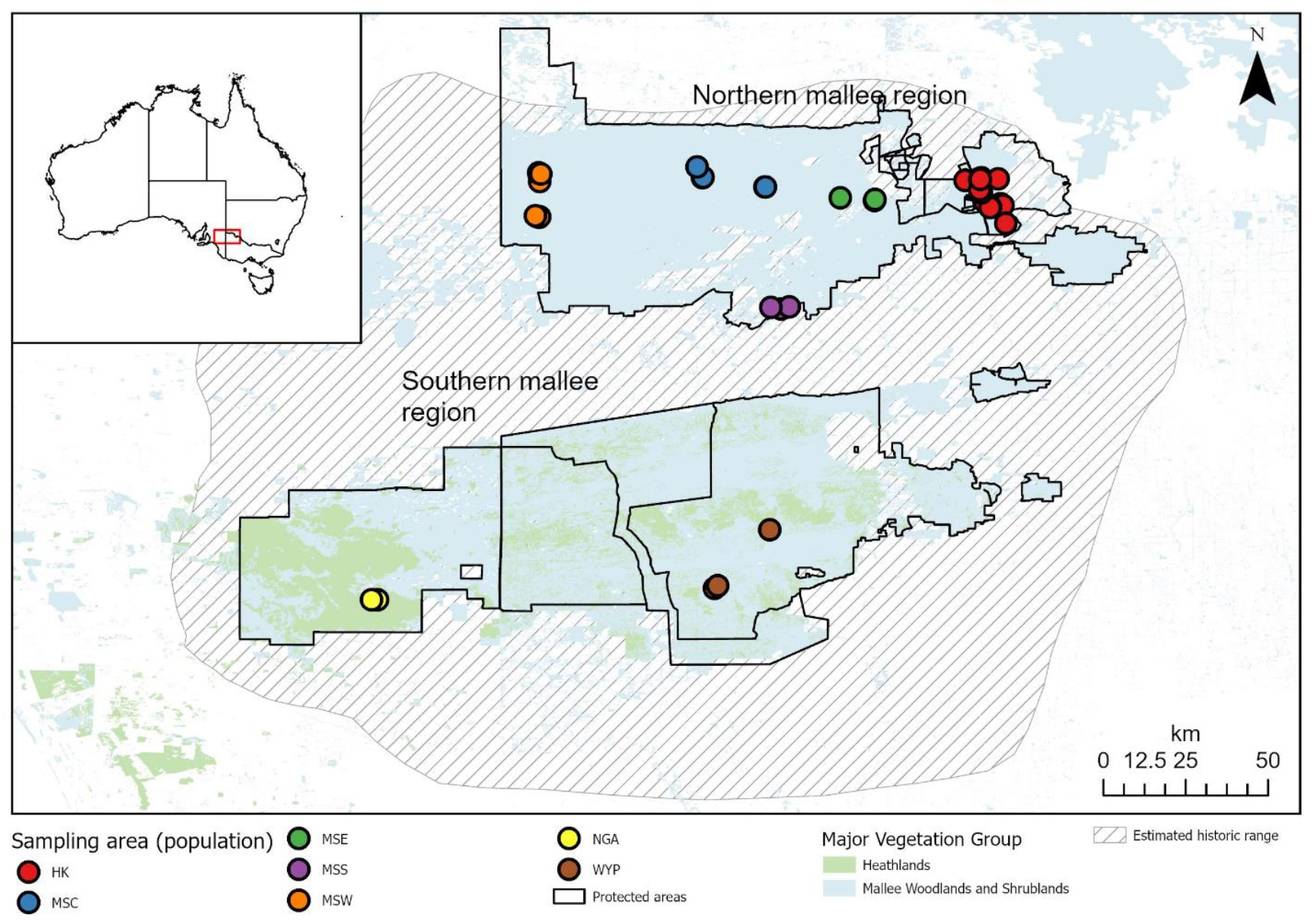
Collection points (coloured circles) of mallee emu-wren genetic material, grouped by sampling area (Table 1). Extant cover of ‘Major Vegetation Classes’ (Australian Government Department of Climate Change Energy the Environment and Water 2023) known to support mallee emu-wren habitat are shaded in blue (Mallee woodlands and shrublands) and green (Heathlands). The actual suitability of local habitat within these broad vegetation classes varies in time depending on time since fire, landscape features, and vegetation composition (Verdon *et al*. 2019, Mitchell *et al*. 2025). Estimated historical range of the mallee emu-wren is represented by hatched area (Higgins *et al*. 2001). A key for sampling area abbreviations is available in table 1. Mallee emu-wrens were extirpated from NGA in 2014.

### Sample collection

In this study we used 72 DNA samples held within Museums Victoria’s biobank repository, collected between 2006 and 2008 from four areas across Murray Sunset National Park, one area in Hattah-Kulkyne National Park, and one in Ngarkat Conservation Park (Brown *et al*. 2013) and three new samples from an isolated mallee emu-wren population collected in Wyperfeld National Park (Verdon *et al*. 2025) in September 2019 using the approach outlined by Mitchell *et al*. (2021). These samples span both continuous tracts of remnant vegetation inhabited by mallee emu-wrens (northern mallee and southern mallee regions on Fig. 1), including six sampling areas currently occupied by mallee emu-wrens and one sampling area from which the species has been locally extirpated (Ngarkat, southern mallee region, Table 1, Fig. 1). Up to 24 µl of blood was drawn from the brachial vein of three newly-collected birds using a heparinised capillary tube and transferred to ethanol. DNA was extracted from these samples using Qiagen’s DNeasy Blood and Tissue Kits following the manufacturer’s instructions.

**Table 1.** The number of genotyped samples obtained from each sampling area (excluding one sample that contained insufficient data for analyses). Extant mallee emu-wren range encompasses two continuous tracts of remnant vegetation, here referred to as northern mallee and southern mallee regions. Hattah-Kulkyne includes sites in Hattah-Kulkyne National Park and Nowingi State Forest. Murray Subset National Park includes four sampling areas (central, east, south, west). Wyperfeld - Wyperfeld National Park. Ngarkat-Ngarkat Conservation Park

| Sampling area abbreviation | Sampling area name | Region | No. of individuals (Female:Male) | Status | Sampling date | Max. distance between individuals |
| --- | --- | --- | --- | --- | --- | --- |
| HK | Hattah-Kulkyne | Northern mallee region | 31 (11:20) | Extant | 2006-2008 | 23.1 km |
| MSW | Murray-Sunset west | Northern mallee region | 11 (5:6) | Extant | 2006-2008 | 13.4 km |
| MSC | Murray-Sunset central | Northern mallee region | 8 (4:4) | Extant | 2006-2008 | 18.1 km |
| MSE | Murray-Sunset east | Northern mallee region | 9 (5:4) | Extant | 2006-2008 | 8.7 km |
| MSS | Murray-Sunset south | Northern mallee region | 5 (2:3) | Extant | 2006-2008 | 4.6 km |
| WYP | Wyperfeld | Southern mallee | 3 (0:3) | Extant | 2019 | 22.4 km |
| NGA | Ngarkat | Southern mallee | 4 (3:1) | **Extinct** | 2006-2008 | 1.5 km |

### Genotyping and data filtering

Genome-wide genotyping of all 75 samples was conducted commercially at Diversity Arrays Technology (Pty Ltd) using their DArTseq™ platform (Kilian *et al*. 2012). Genomic data were generated for 72 individuals comprising 27,727 codominant bi-allelic SNPs: three museum samples contained insufficient DNA for sequencing.

Filtering of genomic data was conducted using the R package *DartR* 2.9.9.5 (Gruber *et al*. 2018). We removed loci that were not 100% reproducible in technical replicates (done for 25% of all samples). Because missing data may be a source of considerable bias during analyses of large genomic datasets (Yi and Latch 2021; Kardos and Waples 2024), we removed all loci with data missing in more than 5% of individuals. One individual with >15% of data missing was removed. We retained only one SNP per sequenced fragment. Finally, we filtered out loci with patterns of genotypes indicative of sex-linkage using the function gl.drop.sexlinked in the package *dartR*.*sexlinked* 1.0.5 (Robledo-Ruiz *et al*. 2025) and phenotypically-determined sex of each individual. Our final filtered autosomal data set comprised 71 individuals from seven sampling areas and included 10,840 binary SNPs with overall 0.69% missing data.

### Genetic analyses

Population genetic structure was estimated using the snmf function in the R package *LEA* 3.20.0 (max k = 10, alpha = 10, tolerance = 10-5, no. iterations = 200, Frichot *et al*. 2015). This function allows the optimum number of clusters (k) to be estimated by generating cross-entropy scores for each potential value of k. Principal coordinate analysis (PCoA) was also used to estimates genetic structure, using the function gl.pcoa in the R package *dartR* 2.9.9.5.

We estimated pairwise Weir and Cockerham *F*_ST_ using the R package *StAMPP* 1.6.3 (Weir and Cockerham 1984; Pembleton *et al*. 2013). We used 100 bootstrap replicates, resampling a subset of loci, to calculate statistical significance and 95% confidence intervals. *F*_ST_ is a frequently used measure of genetic differentiation between geographic groups of organisms. However, mean kinship (MK) has been suggested as a preferable choice for characterising genetic variation among groups of conservation concern (Frankham *et al*. 2017). An individual’s mean kinship is the average co-ancestry it shares with every other individual in a population, including itself (Robledo-Ruiz *et al*. 2022). We calculated pairwise kinship between each individual in the dataset using the beta.dosage function in the R package *hierfstat* 0.5-11 (Goudet 2005). We then averaged these values for all pairwise comparisons between each population to calculate MK between sampling areas. Positive kinship values represent a greater degree of relatedness between a pair of individuals compared with the global average, while negative values represent a lower level of relatedness compared with the global average.

We used the R package *hierfstat* 0.5-11 to calculate observed (*H*_o_) and expected (*H*_s_) heterozygosity, allelic richness (AR) and the fixation index *F*_IS_ for each population, to test for deviations from Hardy Weinberg equilibrium. For each individual, we used the Lynch-Ritland estimator to calculate the inbreeding coefficient *F* using *Coancestry* V1.0.2.0 (Lynch and Ritland 1999; Wang 2011).

We used the R package *strataG* 2.5.5 (Archer *et al*. 2017) to estimate effective population size (Ne) using the linkage-disequilibrium (LDNe) method (Waples *et al*. 2016). We excluded Ngarkat from this analysis, because the population has become extinct, and Wyperfeld because sample sizes <30 may cause significant bias when estimating Ne using the LD method and because WYP is geographically isolated from all other areas supporting mallee emu-wrens (Waples *et al*. 2016). Admixture and PCoA analyses suggested moderate structure between Hattah-Kulkyne and sampling areas in Murray Sunset. Admixture, PCoA, pairwise *F*_ST_ and MK suggested little evidence of structure or differentiation between Murray Sunset central, south or west. Accordingly, we pooled samples from these three areas for Ne analysis. As a result, Ne was calculated for two subsets of individuals: Hattah-Kulkyne (n = 31) and the above-mentioned Murray Sunset areas combined (central, south and west, n = 24). Estimates of Ne calculated from mixed-age samples (as is the case here) using the linkage disequilibrium method may be downwardly biased by 10 – 50%, thus we corrected them for overlapping generations by multiplying the mean and lower CI by 1.1 and mean and upper CI by 1.5 (Waples 2024). Following recommendations of Frankham *et al*. (2014), we interpreted Ne < 1000 as indication that population is at risk of low adaptive potential.

To understand the contribution of each population to overall genetic diversity of the global population of the species, we used *metapop2* 2.5.3 (Lopez-Cortegano *et al*. 2019) to estimate expected heterozygosity within populations (H_S_), Nei’s genetic distance between populations (D_G_), allelic diversity within populations (A_S_, i.e., the sum of average allelic richness across populations minus one), and allelic diversity between populations (D_A_, i.e., average number of alleles present in a population but absent in another, across all pairs of populations). *Metapop2* then removes each population one at a time and recalculates within-population diversity (H_S_, A_S_), and between-population diversity (D_G_, D_A_), to determine the contribution of each population to overall genetic diversity of the species. In this output, positive values of H_S_ or A_S_ indicate that the loss of that population will result in a loss of genetic diversity for the global population. 95% confidence intervals were generated for all values using 1000 bootstrap replicates.

We also used *metapop2* to estimate the optimum percentage of founders that should be sourced from each population to maximise heterozygosity (H) and allelic richness (K) in a hypothetical newly established population of 100 individuals. Samples from Ngarkat were excluded from all metapop2 analyses, because the population is extinct and cannot contribute to future management.

## Results

### Population structure

A low level of genetic differentiation between birds sampled from different areas of remnant vegetation was supported by principal coordinate and admixture analyses (Figs 2 and 3). The first PCoA axis, explaining 4.7% of variance in the dataset, showed a distinction between Hattah-Kulkyne (the easternmost population of the northern mallee region) and all other populations, except a single individual from Hattah-Kulkyne clustered with other populations. Individuals from the southern mallee region (Ngarkat and Wyperfeld) clustered with individuals from the northern mallee region (Murray Sunset central, east, south and west), despite considerable geographic separation between regions (min ∼20 km of agricultural land considered highly unsuitable for mallee emu-wrens). No other meaningful genetic separation of geographic populations was observed (Fig. 2a, b).

**Figure 2.**
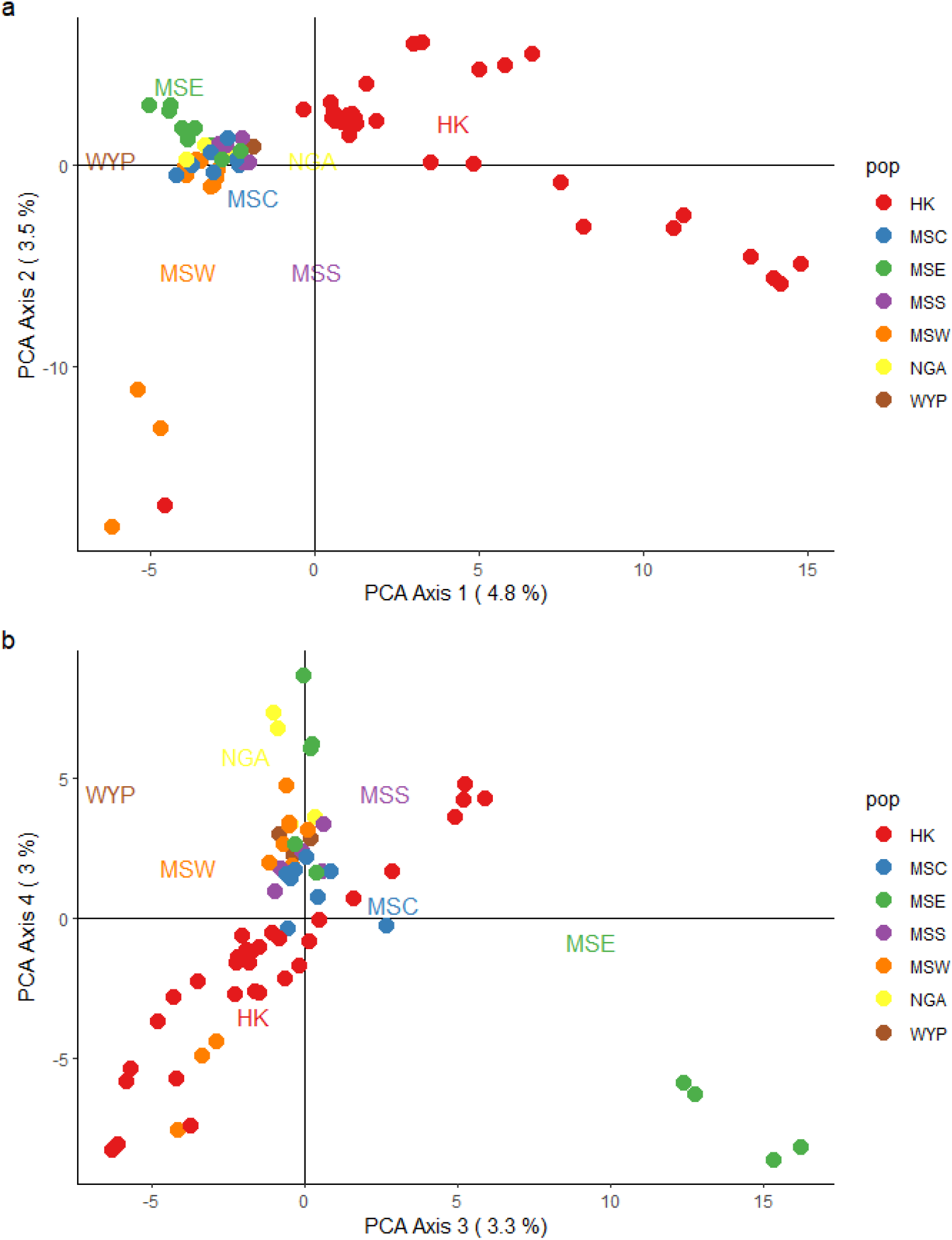
Ordinance plot displaying principal coordinate analysis results for 71 mallee emu-wrens sampled across the contemporary range of the species and one location from which they have become extinct (NGA). a) PCA axes 1 and 2, b) PCA axes 3 and 4. HK = Hattah-Kulkyne, MSC = Murray Sunset central, MSE = Murray Sunset east, MSS = Murray Sunset south, MSW = Murray Sunset west, NGA = Ngarkat, WYP = Wyperfeld.

**Figure 3.**
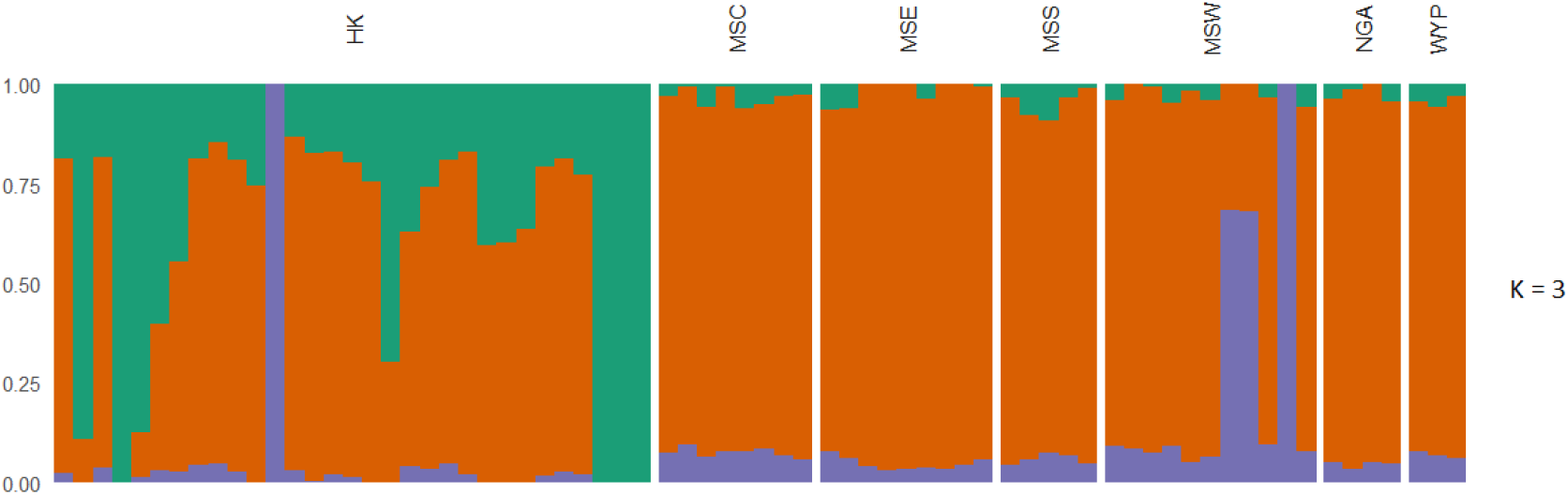
Bar plot displaying results of k=3 structure across 71 mallee emu-wrens sampled across the contemporary range of the species. The optimal number of clusters was determined using the elbow method on cross-entropy scores, generated using the LEA package in R. HK = Hattah-Kulkyne, MSC = Murray Sunset central, MSE = Murray Sunset east, MSS = Murray Sunset south, MSW = Murray Sunset west, NGA = Ngarkat, WYP = Wyperfeld.

Analysis of admixture also showed population structure between Hattah-Kulkyne and all other sampling areas (Fig. 3). The optimal value of k (indicating the number of genetically distinct groups within a sample) for admixture analysis was determined to be three using the elbow method on cross-entropy scores. A small number of individuals were identified in admixture analysis as belonging to a unique cluster in comparison to other individuals from the same sampling area (Fig. 3): these outlying individuals were found to have a high degree of relatedness and were sampled within close proximity to one another.

### Pairwise population differentiation and mean kinship

The results of pairwise *F*_ST_ and MK broadly aligned (Fig. 4), with some exceptions. Relative to other pairwise comparisons, a higher level of differentiation (*F*_ST_ > 0.053, MK < -0.029) was found between Ngarkat and most sampling areas in the northern mallee region (Fig. 4), the only exception being MK between Ngarkat and Murray Sunset west (MK = -0.05). Despite both sampling areas being in the southern mallee region, relatively high differentiation was found between (currently extinct) Ngarkat and Wyperfeld (*F*_ST_ = 0.039, MK = -0.022, Fig. 4). Wyperfeld (southern mallee) had a moderate-high level of differentiation from all sampling areas of the norther mallee region (*F*_*ST*_: 0.031 - 0.05), and low co-ancestry with all sampling areas (MK < -0.029) except with Murray Sunset west (MK = 0.003). Within Murray Sunset areas (northern mallee region), pairs of sampling areas typically had low - moderate differentiation (*F*_ST_: 0.02 – 0.048, MK: -0.022 – 0.19). Murray Sunset east had the greatest level of differentiation from other sampling areas (*F*_ST_: 0.035 – 0.048, MK: -0.022 – 0.009) while Murray Sunset west typically had the greatest degree of co-ancestry (*F*_ST_: 0.025 – 0.048, MK = 0.009 – 0.19). *F*_ST_ and MK indicated a moderate-high level of differentiation between Hattah-Kulkyne and all four Murray Sunset sampling areas (*F*_ST_ = 0.031 – 0.051) and moderate levels of co-ancestry between Hattah-Kulkyne and all Murray Sunset sampling areas (MK = -0.022 – -0.014) except Murray Sunset west, with which MK was relatively high at 0.017 (Fig. 4).

**Figure 4.**
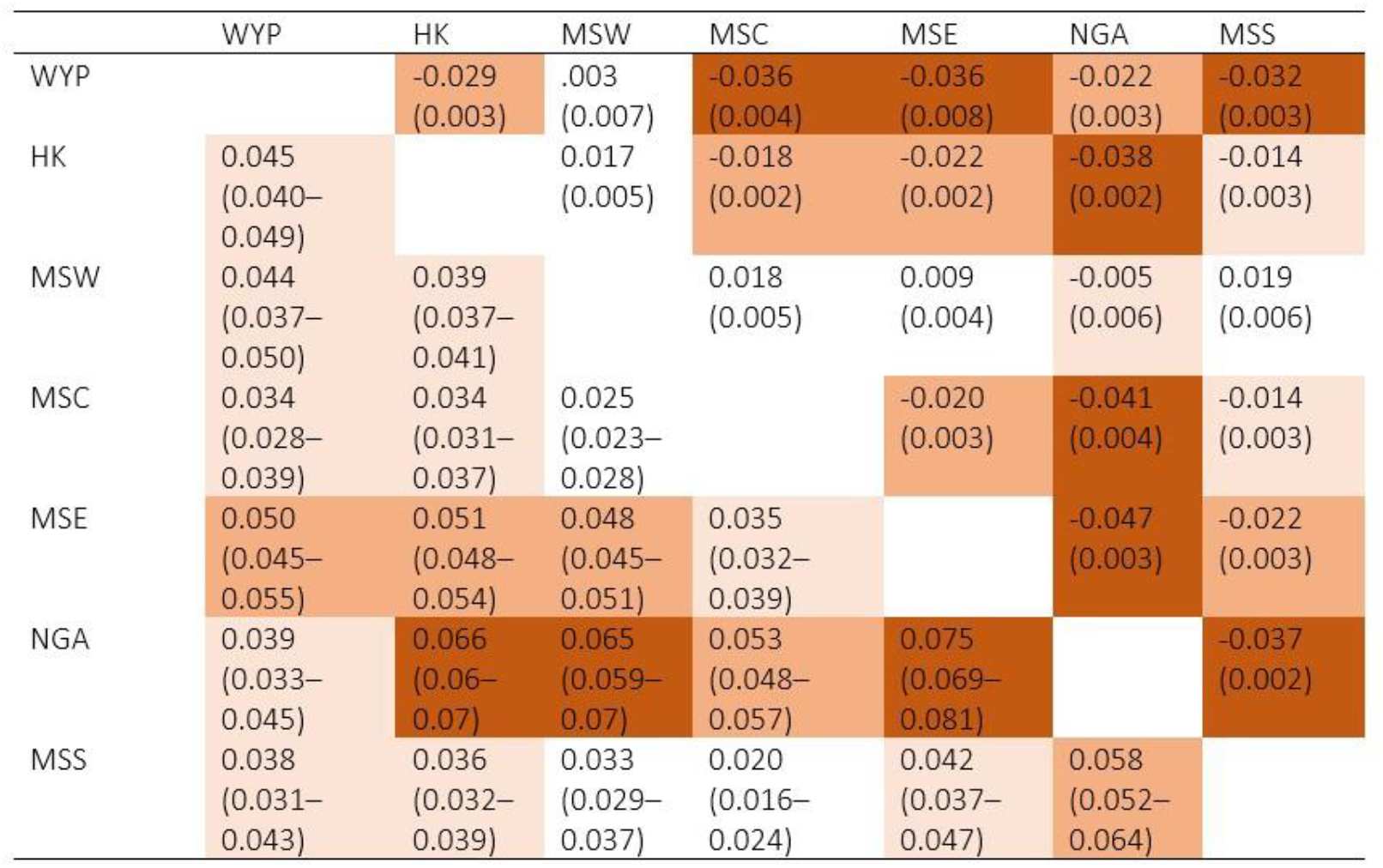
Genetic differentiation between sampling areas. Weir and Cockerham pairwise *F*_ST_ with bootstrap estimates of 95% CI (bottom left semi-matrix) and mean kinship with 2*SE (top right semi-matrix). Mean kinship is scaled such that 0 = global average. All *F*_ST_ values were significant (P value <0.001). colour ramp = level of differentiation between sampling areas. 0 – 25% of range = white, 25-50% of range = light shading, 50-75% of range = moderate shading, 75-100% of range = dark shading.

### Genetic diversity and effective popualtion size

Heterozygosity estimates were similar across all sampling areas (*H*_O_: 0.153 – 0.168, *H*_S_: 0.169 – 0.184, Table 2), as were estimates of allelic richness (AR: 1.252 – 1.273, Table 2). The two southern mallee areas (Ngarkat and Wyperfeld) had considerably lower *F*_IS_ (0.015 – 0.030) than all other sampling areas (0.050 – 0.075). The distribution of individual inbreeding coefficient values did not significantly differ between sampling areas (Figure 5).

**Table 2.** Observed heterozygosity (*H*_O_), expected heterozygosity (*H*_S_), *F*_IS_ and allelic richness (AR) for 71 mallee emu-wrens from six sampling areas across the contemporary range of the species and one location from which mallee emu-wrens have become extinct.

| Sampling area | $H_o$ | $H_s$ | $F_{IS}$ | AR |
| --- | --- | --- | --- | --- |
| Hattah-Kulkyne (n = 31) | 0.162 | 0.178 | 0.075 | 1.266 |
| Murray Sunset central (n = 8) | 0.167 | 0.183 | 0.064 | 1.273 |
| Murray Sunset east (n = 9) | 0.160 | 0.177 | 0.067 | 1.264 |
| Murray Sunset south (n = 5) | 0.165 | 0.181 | 0.050 | 1.269 |
| Murray Sunset west (n = 11) | 0.153 | 0.169 | 0.068 | 1.252 |
| Ngarkat (n = 4) | 0.168 | 0.178 | 0.015 | 1.264 |
| Wyperfeld (n = 3) | 0.167 | 0.184 | 0.030 | 1.272 |

**Figure 5.**
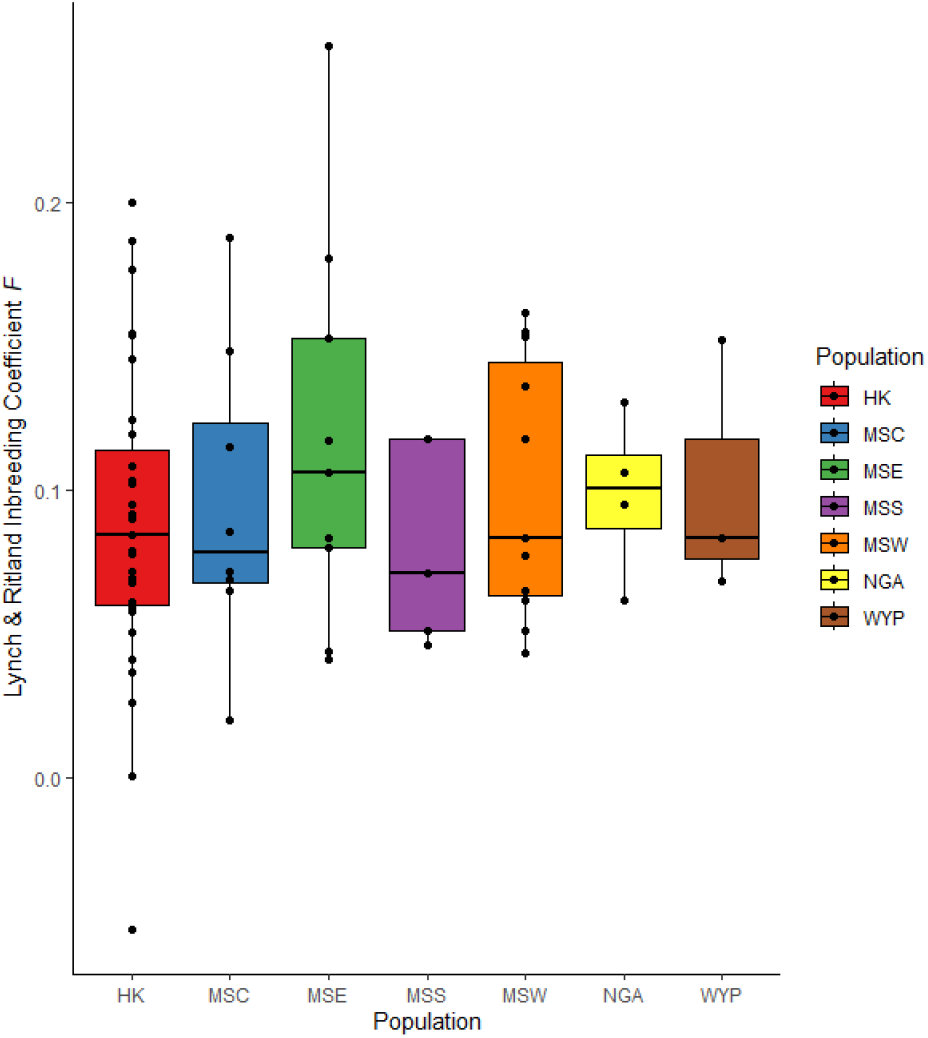
Distributions of individual inbreeding coefficients (Lynch-Ritland *F*), plotted by sampling area. Raw values are plotted in black. HK = Hattah-Kulkyne, MSC = Murray Sunset central, MSE = Murray Sunset east, MSS = Murray Sunset south, MSW = Murray Sunset west, NGA = Ngarkat, WYP = Wyperfeld.

Effective population size (Ne) per generation was estimated to be 47.65 (95% CI: 47.49 - 47.82) for Hattah-Kulkyne, and 87.17 (95% CI: 86.54 – 87.82) for pooled samples from central, southern and western Murray Sunset sampling areas. After applying the adjustments recommended by (Waples 2024) to account for bias caused by generational overlap and mixed age individuals, Ne_adj_ estimated to be to be 52.4 – 71.5 for Hattah-Kulkyne and 95.9 – 130.8 for Murray Sunset central, south and west combined.

### Metapop2 simulations

*Metapop2* revealed considerable differences in the relative contributions of each population to genetic and allelic diversity (Fig. 6). The loss of Hattah-Kulkyne would result in the greatest loss of global within-population heterozygosity (H_S_), followed by Murray Sunset central, whereas loss of any one other area would not decrease heterozygosity of a total meta-population. Meanwhile, the loss of any one of Hattah-Kulkyne, Murray Sunset central, Murray Sunset south or Wyperfeld would result in a loss of allelic diversity (A_S_) for the global mallee emu-wren meta-population. Murray Sunset west was the only population that did not contribute unique genetic (H_S_) or allelic (A_S_) diversity to the global meta-population (Fig. 6).

**Figure 6.**
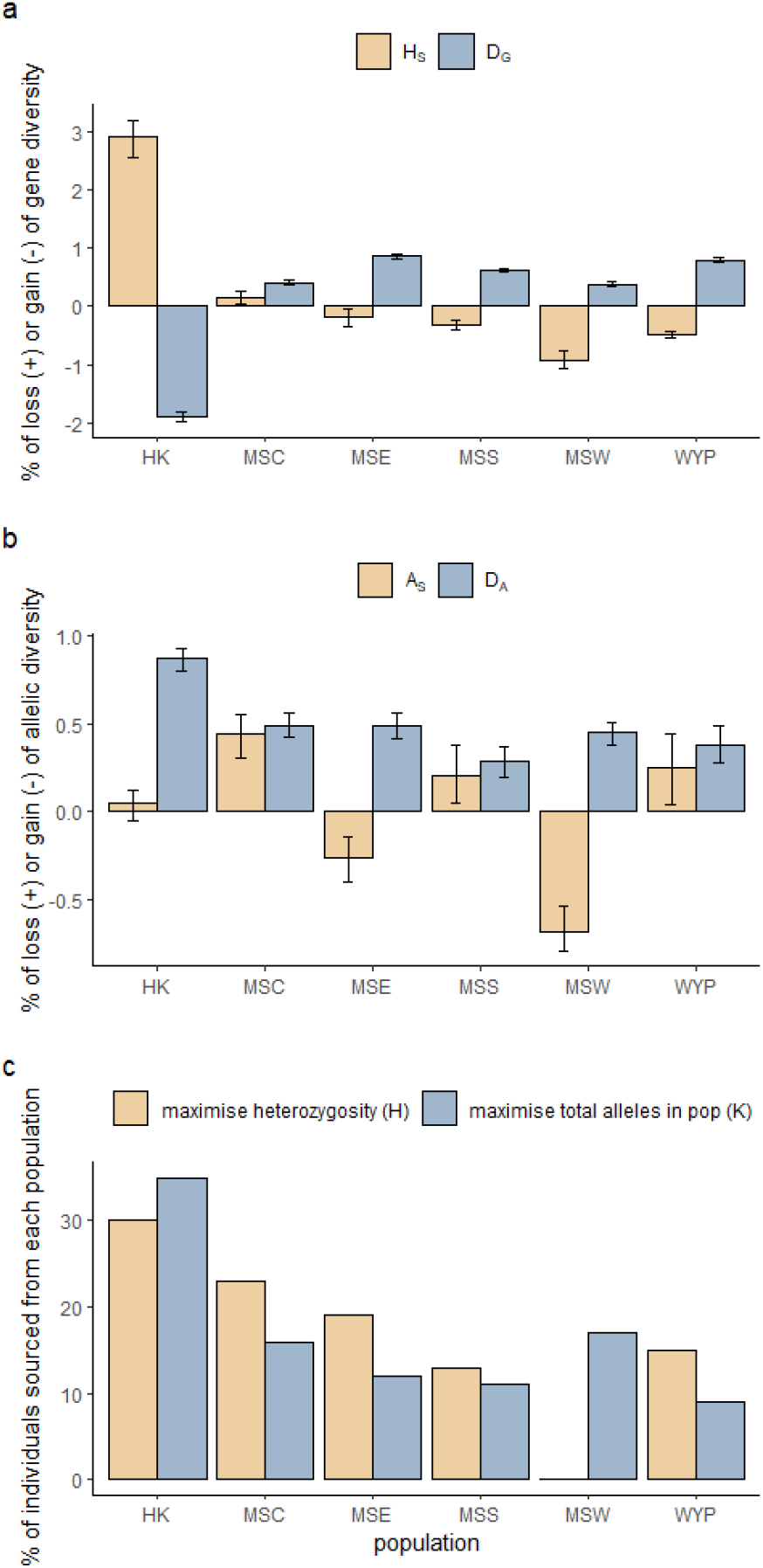
**a)** The contribution of each population to genetic diversity within populations (expected heterozygosity H_S_) and between populations (Nei’s minimum genetic distance D_G_). b) The contribution of each population to allelic diversity within population (A_S_) and between populations (D_A_). For panels a and b, positive values represent the percentage of loss while negative values represent the level of gain in diversity after removal of each population from the global mallee emu-wren population. Error bars show 95% bootstrap limits for each estimate. c) percentage of individuals from each population needed in a synthetic pool of 100 individuals (e.g., a hypothetical translocation to establish a new population) to maximise expected heterozygosity (H) or total number of alleles (K).

Sourcing founders from all 6 extant populations would be required to maximise allelic richness (K) in a newly established translocated population of mallee emu-wrens, with the greatest contribution coming from Hattah Kulkyne (Fig. 6). Maximum heterozygosity (H) could be achieved by sourcing founders from all populations with the exception of MSW, with the greatest contribution, again, coming from Hattah-Kulkyne (Fig. 6).

## Discussion

The global mallee emu-wren meta-population is likely to have been historically characterised by spatially and temporally variable connectivity and gene flow (Brown *et al*. 2013). However, habitat loss and fragmentation, exacerbated by inappropriate fire regimes and drought have disrupted meta-population dynamics, leading to isolated sub-populations and, in many cases, local extinctions. We assessed genetic diversity, differentiation and structure across the global mallee emu-wren population using >10,000 high-quality SNPs from 71 individuals across six sampling areas within the contemporary range of the species and one from which they have become extinct. We found weak genetic structure among mallee emu-wren sampling areas, notably between Hattah-Kulkyne and all other areas. Limited differentiation across the species’ range supports the conclusion of Brown *et al*. (2013), based on a less informative data set, that the global mallee emu-wren population may be considered a single genetic unit for management purposes. Effective population size estimates for both the Hattah-Kulkyne and pooled sample of Murray Sunset central, south and west were <1000, indicating low adaptive potential (Frankham *et al*. 2014). To mitigate inbreeding depression within sub-populations and increase adaptive potential across the global population of the species, we recommend gene-pool mixing between extant mallee emu-wren sub-populations. Likewise, we recommend that founders for any newly established mallee emu-wren population should be sourced from all available sub-populations that are sufficiently demographically robust to sustain harvest of founders, with particular focus on combining Hattah-Kulkyne and other sources.

Hattah-Kulkyne had the greatest difference from all other sampling areas, despite close proximity to other sampling areas within the northern mallee region. Little structure was evident among three of the four sampling locations in Murray Sunset National Park (central, southern and western). Surprisingly, these sampling areas were also not distinct from southern mallee sampling areas (Ngarkat and Wyperfeld). This result contrasts that of Brown *et al*. (2013) whose TESS analysis incorporating spatial data revealed Ngarkat (a single location of the southern mallee region in that study) belonging to one genetic cluster, whereas all other sampled areas excluding eastern Murray Sunset belonged to an alternative cluster. The genetic distinction of Hattah-Kulkyne may be due to restricted dispersal through the ∼400 km^2^ Raak Plain, which lies between Murray Sunset and Hattah-Kulkyne. Higher salinity of this area creates conditions not suitable for *Triodia* growth and, consequently, Raak Plain is likely to have been a barrier for mallee emu-wrens for at least the last several thousand years. Since the early 20^th^-century, gene flow among mallee emu-wrens in Murray Sunset and Hattah-Kulkyne has been feasible only through an ∼5 km wide strip of native vegetation connecting the two reserves, south of Raak Plain. One individual sampled in Hattah-Kulkyne (which appears as an obvious outlier in figures 2 and 3), was found to have a high degree of co-ancestry with individuals sampled in Murray Sunset west. This may represent a rare case of dispersal between these geographically distant sampling areas, though the possibility of a sampling error must also be considered.

The low level of genetic difference between the southern and northern mallee regions is difficult to explain. These regions have been separated by a broad expanse of agricultural land since at least the 1950s (Clarke *et al*. 2001), and the poor dispersal capability of mallee emu-wrens across agricultural land would argue against the occurrence of gene flow between these regions in the last >80 years. Prior to widespread land clearing, the intervening landscape would have been characterised by a matrix of Woorinen sands mallee and loamy sands mallee that cycled through phases of suitability for mallee emu-wrens depending on prevailing climate and fire history (Brown *et al*. 2013; Department of the Environment 2014). The lack of structure evident in this study could be due to a relatively low level of genetic drift in the decades since separation between the southern and northern mallee regions, though this is unlikely based on life history and the small population size of mallee emu-wrens in the southern mallee region. It is more likely that regular fire-driven bottlenecks and strong genetic drift have reduced the species’ historical genetic diversity to the point that only common alleles are present in the meta-population, precluding detection of emerging population structure through retention of different sets of rare alleles. Calculating unbiased genetic diversity using a reference genome and genome-wide sequencing data (Pavlova *et al*. 2025) would enable comparison of mallee emu-wren diversity with other threatened species. Increasing sample sizes from the southern region might increase the power to detect genetic differentiation across regions.

Heterozygosity and allelic richness differed little between mallee emu-wren populations. This contrasted with results presented by (Brown *et al*. 2013), who found comparable levels of heterozygosity between sampling areas but lower levels of allelic richness in Ngarkat and Murray Sunset East compared with other sampling areas. Observed heterozygosity at all sampling areas was lower than expected heterozygosity: very high co-ancestry between some individuals within sampling locations suggests sampling of family groups contributed to localized population genetic structure (i.e. Wahlund effect, Wahlund 1928).

Low Ne <100 estimates from this study for Hattah-Kulkyne, and three Murray Sunset areas (central, south and west) present a clear trigger for genetic augmentation to mitigate loss of fitness and adaptive potential (Frankham *et al*. 2017; Ralls *et al*. 2018). The results of metapop2 show that Hattah-Kulkyne and Murray Sunset central are contributing unique genetic diversity to the global mallee emu-wren population, highlighting the potential benefit of using these areas as a source of genetic augmentation to mitigate inbreeding depression in other sampling areas. The results of metapop2 also indicate that gene-pool mixing among all sampling areas except Murray Sunset east and west have potential to improve allelic diversity, thus providing greater adaptive potential, in recipient sub-populations. Frankham *et al*. (2017) provide clear guidelines for the number of migrants required to prevent damaging inbreeding, recommending five or more effective migrants (a migrant that successfully contributed its genes to a local population) per generation (and more for populations that are already inbred). Because Ne is, on average, ∼13% of N, Frankham *et al*. (2017)’s recommendations translate to ∼38 migrants per generation. Given small population sizes, short generation time, and the inherent risk of capture, transport and release (Dickens *et al*. 2010; Berger-Tal *et al*. 2019), sustaining such levels of reciprocal migration between mallee emu-wren sub-populations is likely to be demographically risky (Mitchell *et al*. 2021). On top of this, sufficient resources to implement such a strategy are not currently available. Despite these challenges, the risk of a ‘do nothing’ approach is significant and Frankham *et al*. (2017) highlight that when a recipient population is inbred, any level of immigration is better than none. Genetic augmentation at a small scale would provide some mitigation of inbreeding depression and improvement in adaptive potential, while providing an opportunity for greater genetic sampling and research among extant mallee emu-wren populations.

To maximise allelic richness in future translocated populations, founders should be drawn from all extant populations of mallee emu-wrens. Conservation managers must ensure that removing individuals from wild populations for translocation is sustainable (Mitchell *et al*. 2022). Little is known about mallee emu-wren population demographics, particularly in Wyperfeld. Given demographic uncertainty and potential risk, populations such as Murray Sunset west found to contribute little unique allelic diversity to the global mallee emu-wren population, should still be considered as sources for translocation given shared diversity, robust population size and levels of population connectivity suggesting greater resilience to the impacts of translocation harvest. Of 78 mallee emu-wrens reintroduced to Ngarkat in 2018, at least 15 were confirmed to have contributed to a nesting attempt (based on discoveries of nests containing eggs/chicks or the presence of fledglings with adults, Mitchell *et al*. 2021). The ratio is more favourable when considering only those birds released in spring compared with autumn (at least 12 of 38 birds). The demographic impact of translocation on source populations may also be reduced by aligning harvest with favourable climatic conditions (Verdon *et al*. 2021). Careful management should allow gene-pool mixing between mallee emu-wren populations while alleviating many of the risks of harvesting individuals for translocation.

### Conclusion

Providing best-practice genetic management to the global mallee emu-wren meta-population will provide the greatest probability of ensuring its long-term conservation by maximising fitness and adaptive potential and ensuring that any newly established translocated populations have the greatest probability of persistence. Notwithstanding considerable current logistical and resource constraints, adopting an informed best-practice genetic management approach is hampered by small sample size, particularly in the southern mallee, and the time elapsed since many of the available genetic samples were taken. Despite these limitations, we have demonstrated that mallee emu-wren sub-populations have effective population sizes indicative of dangerous inbreeding depression, including in areas considered ‘hotspots’ for this Endangered species. We recommend genetic augmentation as an urgent strategy to mitigate inbreeding depression and improve adaptive potential across the global mallee emu-wren meta-population. Genetic augmentation will also provide an opportunity for increasing the amount of genetic data available to guide conservation management of the mallee emu-wren.

## Acknowledgements

This research was undertaken under Department of Environment, Land, Water and Planning Research Permit no. 10009048 and Monash University Ethics Approval no. 18349. We are grateful to Sarah Brown who originally collected most of the samples used in this study and to Joanna Sumner and Museums Victoria for providing access to those samples. Finally, we would like to thank Owen Lishmund, Ben Viola and Michael Roast who volunteered their time to help catch mallee emu-wrens.

## Data Availability Statement

Upon publication, the dataset used in this study will be published on OPAL, La Trobe University’s open access institutional repository.

## References

Amos N, Harrisson KA, Radford JQ, White M, Newell G, Mac Nally R, Sunnucks P, Pavlova A (2014) Species- and sex-specific connectivity effects of habitat fragmentation in a suite of woodland birds. Ecology 95(6), 1556–1568. doi:10.1890/13-1328.1

Andersen A, Simcox DJ, Thomas JA, Nash DR (2014) Assessing reintroduction schemes by comparing genetic diversity of reintroduced and source populations: A case study of the globally threatened large blue butterfly (Maculinea anion). Biological Conservation 175, 34–41. doi:10.1016/j.biocon.2014.04.009

Archer FI, Adams PE, Schneiders BB (2017) stratag: An r package for manipulating, summarizing and analysing population genetic data. Molecular Ecology Resources 17(1), 5–11. doi:10.1111/1755-0998.12559

Australian Government Department of Climate Change Energy the Environment and Water (2023) National Vegetation Information System (NVIS) Version 6. [Verified 01/06/2024]

Berger-Tal O, Blumstein DT, Swaisgood RR (2019) Conservation translocations: a review of common difficulties and promising directions. Animal Conservation 23(2), 121–131. doi:10.1111/acv.12534

Bertassello LE, Bertuzzo E, Botter G, Jawitz JW, Aubeneau AF, Hoverman JT, Rinaldo A, Rao PSC (2021) Dynamic spatio-temporal patterns of metapopulation occupancy in patchy habitats. Royal Society Open Science 8(1), 201309. doi:10.1098/rsos.201309. PMCID: PMC7890491

Brown S, Clarke M, Clarke R (2009) Fire is a key element in the landscape-scale habitat requirements and global population status of a threatened bird: The Mallee Emu-wren (Stipiturus mallee). Biological Conservation 142, 432–445. doi:10.1016/j.biocon.2008.11.005

Brown SM, Harrisson KA, Clarke RH, Bennett AF, Sunnucks P (2013) Limited population structure, genetic drift and bottlenecks characterise an endangered bird species in a dynamic, fire-prone ecosystem. PLoS One 8(4), e59732. doi:10.1371/journal.pone.0059732. PMCID: PMC3634030

Camak DT, Osborne MJ, Turner TF (2021) Population genomics and conservation of Gila Trout (Oncorhynchus gilae). Conservation Genetics 22(5), 729–743. doi:10.1007/s10592-021-01355-0

Ceballos G, Ehrlich PR, Barnosky AD, García A, Pringle RM, Palmer TM (2015) Accelerated modern human–induced species losses: Entering the sixth mass extinction. Science advances 1(5), e1400253.

Clarke RH, Gordon IR, Clarke MF (2001) Intraspecific phenotypic variability in the black-eared miner (Manorina melanotis); human-facilitated introgression and the consequences for an endangered taxon. Biological Conservation 99(2), 145–155.

Connell J, Hall MA, Nimmo DG, Watson SJ, Clarke MF, Parr C (2022) Fire, drought and flooding rains: The effect of climatic extremes on bird species’ responses to time since fire. Diversity and Distributions 28, 417–438. doi:10.1111/ddi.13287

Cook CN, Sgro CM (2017) Aligning science and policy to achieve evolutionarily enlightened conservation. Conservation Biology 31(3), 501–512. doi:10.1111/cobi.12863

Courchamp F, Berec L, Gascoigne J (2008) ‘Allee effects in ecology and conservation.’ (Oxford University Press: New York)

Department of the Environment (2014) Estimated Pre-1750 Major Vegetation Subgroups. Bioregional Assessment Source Dataset. (Bioregional Assessment Source Dataset) Available at http://data.bioregionalassessments.gov.au/dataset/2208babe-8e88-4423-91e6-b9a3fa6f31b6

Dickens MJ, Delehanty DJ, Michael Romero L (2010) Stress: An inevitable component of animal translocation. Biological Conservation 143(6), 1329–1341. doi:10.1016/j.biocon.2010.02.032

Feuerstein CA, Kovach RP, Kruse CG, Jaeger ME, Bell DA, Robinson ZL, Whiteley AR (2024) Genetic variation and hybridization determine the outcomes of conservation reintroductions. Conservation Letters 17(5). doi:10.1111/conl.13049

Frankham R (2015) Genetic rescue of small inbred populations: meta-analysis reveals large and consistent benefits of gene flow. Mol Ecol 24(11), 2610–2618. doi:10.1111/mec.13139

Frankham R, Ballou JD, Ralls K, Eldridge M, Dudash MR, Fenster CB, Lacy RC, Sunnucks P (2017) ‘Genetic Management of Fragmented Animal and Plant Populations.’ 10.1093/oso/9780198783398.001.0001

Frankham R, Bradshaw CJA, Brook BW (2014) Genetics in conservation management: Revised recommendations for the 50/500 rules, Red List criteria and population viability analyses. Biological Conservation 170, 56–63. doi:10.1016/j.biocon.2013.12.036

Frichot E, François O, O’Meara B (2015) LEA: An R package for landscape and ecological association studies. Methods in Ecology and Evolution 6(8), 925–929. doi:10.1111/2041-210x.12382

Goudet J (2005) Hierfstat, a package for R to compute and test hierarchical F-statistics. Molecular Ecology Notes 5(1), 184–186.

Gruber B, Unmack PJ, Berry OF, Georges A (2018) dartr: An r package to facilitate analysis of SNP data generated from reduced representation genome sequencing. Molecular Ecology Resources 18(3), 691–699.

Grueber CE, Sutton JT, Heber S, Briskie JV, Jamieson IG, Robertson BC (2017) Reciprocal translocation of small numbers of inbred individuals rescues immunogenetic diversity. Molecular Ecology 26(10), 2660–2673. doi:10.1111/mec.14063

Hanski I (1998) Metapopulation dynamics. Nature 396, 41–49.

He X, Johansson ML, Heath DD (2016) Role of genomics and transcriptomics in selection of reintroduction source populations. Conserv Biol 30(5), 1010–1018. doi:10.1111/cobi.12674

Heber S, Varsani A, Kuhn S, Girg A, Kempenaers B, Briskie J (2013) The genetic rescue of two bottlenecked South Island robin populations using translocations of inbred donors. Proceedings of the Royal Society B: Biological Sciences 280(1752), 20122228. doi:10.1098/rspb.2012.2228. PMCID: PMC3574298

Higgins PJ, Peter JM, Steele WK (2001) ‘Handbook of Australian, New Zealand and Antarctic Birds. Volume 5.’ (Oxford University Press: Melbourne)

Hoban S, Bruford MW, Funk WC, Galbusera P, Griffith MP, Grueber CE, Heuertz M, Hunter ME, Hvilsom C, Stroil BK, Kershaw F, Khoury CK, Laikre L, Lopes-Fernandes M, MacDonald AJ, Mergeay J, Meek M, Mittan C, Mukassabi TA, O’Brien D, Ogden R, Palma-Silva C, Ramakrishnan U, Segelbacher G, Shaw RE, Sjogren-Gulve P, Velickovic N, Vernesi C (2021) Global Commitments to Conserving and Monitoring Genetic Diversity Are Now Necessary and Feasible. Bioscience 71(9), 964–976. doi:10.1093/biosci/biab054. PMCID: PMC8407967

Hundertmark KJ, Van Daele LJ (2010) Founder effect and bottleneck signatures in an introduced, insular population of elk. Conservation Genetics 11(1), 139–147. doi:10.1007/s10592-009-0013-z

Hunt T, Mitchell W, Boulton R, Hedger C, Ireland L (2019) Cooperative breeding recorded in the endangered Mallee Emu-wren Stipiturus mallee. Australian Field Ornithology 36, 163–167. doi:10.20938/afo36163167

Indigo N, Boulton R, Lau J, Fullagar A, Howling G (2023) Threatened Mallee Birds Conservation Action Plan. (BirdLife Australia)

Kardos M, Waples RS (2024) Low-coverage sequencing and Wahlund effect severely bias estimates of inbreeding, heterozygosity and effective population size in North American wolves. Mol Ecol, e17415. doi:10.1111/mec.17415

Keller LF, Waller DM (2002) Inbreeding effects in wild populations. Trends in Ecology & Evolution 17(5), 230–241.

Kilian A, Wenzl P, Huttner E, Carling J, Xia L, Blois H, Caig V, Heller-Uszynska K, Jaccoud D, Hopper C (2012) Diversity arrays technology: a generic genome profiling technology on open platforms. In ‘Data production and analysis in population genomics’. pp. 67–89. (Springer: New York, NY)

Knipler ML, Gracanin A, Mikac KM (2023) Conservation genomics of an endangered arboreal mammal following the 2019-2020 Australian megafire. Scientific Reports 13(1), 480. doi:10.1038/s41598-023-27587-3. PMCID: PMC9831986

Laikre L, Hoban S, Bruford MW, Segelbacher G, Allendorf FW, Gajardo G, Rodríguez AG, Hedrick PW, Heuertz M, Hohenlohe PA (2020) Post-2020 goals overlook genetic diversity. Science 367(6482), 1083–1085.

Langridge J, Sordello R, Reyjol Y (2021) Existing evidence on the outcomes of wildlife translocations in protected areas: a systematic map. Environmental Evidence 10(1), 29. doi:10.1186/s13750-021-00236-w

Liddell E, Sunnucks P, Cook CN (2021) To mix or not to mix gene pools for threatened species management? Few studies use genetic data to examine the risks of both actions, but failing to do so leads disproportionately to recommendations for separate management. Biological Conservation 256, 109072. doi:10.1016/j.biocon.2021.109072

Lopez-Cortegano E, Perez-Figueroa A, Caballero A (2019) metapop2: Re-implementation of software for the analysis and management of subdivided populations using gene and allelic diversity. Molecular Ecology Resources 19(4), 1095–1100. doi:10.1111/1755-0998.13015

Lutz ML, Tonkin Z, Yen JDL, Johnson G, Ingram BA, Sharley J, Lyon J, Chapple DG, Sunnucks P, Pavlova A (2021) Using multiple sources during reintroduction of a locally extinct population benefits survival and reproduction of an endangered freshwater fish. Evol Appl 14(4), 950–964. doi:10.1111/eva.13173. PMCID: PMC8061264

Lynch M, Ritland K (1999) Estimation of pairwise relatedness with molecular markers. Genetics 152(4), 1753–1766.

Malone EW, Perkin JS, Leckie BM, Kulp MA, Hurt CR, Walker DM (2018) Which species, how many, and from where: Integrating habitat suitability, population genomics, and abundance estimates into species reintroduction planning. Glob Chang Biol 24(8), 3729–3748. doi:10.1111/gcb.14126

Mitchell WF, Boulton RL, Ireland L, Hunt TJ, Verdon SJ, Olds LGM, Hedger C, Clarke RH (2021) Using experimental trials to improve translocation protocols for a cryptic, endangered passerine. Pacific Conservation Biology 28(1), 68–79. doi:10.1071/pc20097

Mitchell WF, Boulton RL, Sunnucks P, Clarke RH (2022) Are we adequately assessing the demographic impacts of harvesting for wild-sourced conservation translocations? Conservation Science and Practice 4(1), e569. doi:10.1111/csp2.569

Mitchell WF, Paton DC, Connell J, Clarke RH, Verdon SJ (2025) Rainfall immediately before and after fire promotes longterm occurrence of a rare, fire-sensitive passerine. In Review.

Nei M, Maruyama T, Chakraborty R (1975) The bottleneck effect and genetic variability in populations. Evolution 29, 1–10.

Newbold T, Hudson LN, Hill SL, Contu S, Lysenko I, Senior RA, Borger L, Bennett DJ, Choimes A, Collen B, Day J, De Palma A, Diaz S, Echeverria-Londono S, Edgar MJ, Feldman A, Garon M, Harrison ML, Alhusseini T, Ingram DJ, Itescu Y, Kattge J, Kemp V, Kirkpatrick L, Kleyer M, Correia DL, Martin CD, Meiri S, Novosolov M, Pan Y, Phillips HR, Purves DW, Robinson A, Simpson J, Tuck SL, Weiher E, White HJ, Ewers RM, Mace GM, Scharlemann JP, Purvis A (2015) Global effects of land use on local terrestrial biodiversity. Nature 520(7545), 45–50. doi:10.1038/nature14324

Nistelberger HM, Roycroft E, Macdonald AJ, McArthur S, White LC, Grady PGS, Pierson J, Sims C, Cowen S, Moseby K, Tuft K, Moritz C, Eldridge MDB, Byrne M, Ottewell K (2025) Genetic mixing in conservation translocations increases diversity of a keystone threatened species, Bettongia lesueur. Mol Ecol 34(17), e17119. doi:10.1111/mec.17119. PMCID: PMC12376963

Pavlova A, Tonkin Z, Pearce L, Robledo-Ruiz DA, Lintermans M, Ingram BA, Lyon J, Beitzel M, Broadhurst B, Rourke ML, Sturgiss F, Lake E, Castrejon-Figueroa J, Stocks JR, Sunnucks P (2025) A Shift to Metapopulation Genetic Management for Persistence of a Species Threatened by Fragmentation: The Case of an Endangered Australian Freshwater Fish. Mol Ecol, e70005. doi:10.1111/mec.70005

Pembleton LW, Cogan NOI, Forster JW (2013) StAMPP: an R package for calculation of genetic differentiation and structure of mixed-ploidy level populations. Molecular Ecology Resources 13(5), 946–952. doi:10.1111/1755-0998.12129

Piana PA, Gubiani ÉA, Gomes LC (2014) A modified metapopulation model to predict colonisation and extinction rates in fragmented aquatic systems. Ecological Engineering 73, 26–30. doi:10.1016/j.ecoleng.2014.09.024

Pierson JC, Coates DJ, Oostermeijer JGB, Beissinger SR, Bragg JG, Sunnucks P, Schumaker NH, Young AG (2016) Genetic factors in threatened species recovery plans on three continents. Frontiers in Ecology and the Environment 14(8), 433–440. doi:10.1002/fee.1323

Ralls K, Ballou JD, Dudash MR, Eldridge MDB, Fenster CB, Lacy RC, Sunnucks P, Frankham R (2018) Call for a Paradigm Shift in the Genetic Management of Fragmented Populations. Conservation Letters 11(2), e12412. doi:10.1111/conl.12412

Rick K, Byrne M, Cameron S, Cooper SJB, Dunlop J, Hill B, Lohr C, Mitchell NJ, Moritz C, Travouillon KJ, von Takach B, Ottewell K (2023) Population genomic diversity and structure in the golden bandicoot: a history of isolation, extirpation, and conservation. Heredity (Edinb) 131(5-6), 374–386. doi:10.1038/s41437-023-00653-2. PMCID: PMC10673901

Robledo-Ruiz DA, Austin L, Amos JN, Castrejon-Figueroa J, Harley DKP, Magrath MJL, Sunnucks P, Pavlova A (2025) Easy-to-use R functions to separate reduced-representation genomic datasets into sex-linked and autosomal loci, and conduct sex assignment. Molecular Ecology Resources 25(5), e13844. doi:10.1111/1755-0998.13844. PMCID: PMC12142712

Robledo-Ruiz DA, Pavlova A, Clarke RH, Magrath MJL, Quin B, Harrisson KA, Gan HM, Low GW, Sunnucks P (2022) A novel framework for evaluating in situ breeding management strategies in endangered populations. Molecular Ecology Resources 22(1), 239–253. doi:10.1111/1755-0998.13476

Saccheri I, Kuussaari M, Kankare M, Vikman P, Fortelius W, Hanski I (1998) Inbreeding and extinction in a butterfly metapopulation. Nature 392(6675), 491–494. doi:10.1038/33136

Schlaepfer DR, Braschler B, Rusterholz HP, Baur B (2018) Genetic effects of anthropogenic habitat fragmentation on remnant animal and plant populations: a meta-analysis. Ecosphere 9(10), e02488.

Scott PA, Allison LJ, Field KJ, Averill-Murray RC, Shaffer HB (2020) Individual heterozygosity predicts translocation success in threatened desert tortoises. Science 370(6520), 1086–1089.

Shaw RE, Farquharson KA, Bruford MW, Coates DJ, Elliott CP, Mergeay J, Ottewell KM, Segelbacher G, Hoban S, Hvilsom C, Perez-Espona S, Rungis D, Aravanopoulos F, Bertola LD, Cotrim H, Cox K, Cubric-Curik V, Ekblom R, Godoy JA, Konopinski MK, Laikre L, Russo IM, Velickovic N, Vergeer P, Vila C, Brajkovic V, Field DL, Goodall-Copestake WP, Hailer F, Hopley T, Zachos FE, Alves PC, Biedrzycka A, Binks RM, Buiteveld J, Buzan E, Byrne M, Huntley B, Iacolina L, Keehnen NLP, Klinga P, Kopatz A, Kurland S, Leonard JA, Manfrin C, Marchesini A, Millar MA, Orozco-terWengel P, Ottenburghs J, Posledovich D, Spencer PB, Tourvas N, Unuk Nahberger T, van Hooft P, Verbylaite R, Vernesi C, Grueber CE (2025) Global meta-analysis shows action is needed to halt genetic diversity loss. Nature 638(8051), 704–710. doi:10.1038/s41586-024-08458-x. PMCID: PMC11839457

Verdon SJ, Makdissi R, Mitchell WF, Boulton RL, Radford JQ (2025) Benefits of Modelling Abundance for Rare Species Conservation: A Case Study With Multiple Birds Across One Million Hectares. Diversity and Distributions 31(1), e13956. doi:10.1111/ddi.13956

Verdon SJ, Mitchell WF, Clarke MF (2021) Can flexible timing of harvest for translocation reduce the impact on fluctuating source populations? Wildlife Research 48, 458–469. doi:10.1071/WR20133

Verdon SJ, Watson SJ, Clarke MF (2019) Modeling variability in the fire response of an endangered bird to improve fire-management. Ecological Applications 29(8), e01980. doi:10.1002/eap.1980

Wahlund S (1928) Composition of populations and correlation appearances viewed in relation to the studies of inheritance. Hereditas 11, 65–106.

Wang J (2011) COANCESTRY: a program for simulating, estimating and analysing relatedness and inbreeding coefficients. Molecular ecology resources 11(1), 141–145.

Waples RK, Larson WA, Waples RS (2016) Estimating contemporary effective population size in non-model species using linkage disequilibrium across thousands of loci. Heredity (Edinb) 117(4), 233–240. doi:10.1038/hdy.2016.60. PMCID: PMC5026758

Waples RS (2024) Practical application of the linkage disequilibrium method for estimating contemporary effective population size: A review. Molecular Ecology Resources 24(1), e13879. doi:10.1111/1755-0998.13879

Weeks AR, Stoklosa J, Hoffmann AA (2016) Conservation of genetic uniqueness of populations may increase extinction likelihood of endangered species: the case of Australian mammals. Frontiers in Zoology 13(1), 1–9.

Weir BS, Cockerham CC (1984) Estimating F-statistics for the analysis of population structure. Evolution 38(6), 1358–1370.

Yi X, Latch EK (2021) Nonrandom missing data can bias Principal Component Analysis inference of population genetic structure. Molecular Ecology Resources 00, 1–10. doi:10.1111/1755-0998.13498

